# Relationship between high-frequency activity in the cortical sensory and the motor hand areas, and their myelin content

**DOI:** 10.1101/2021.12.02.470914

**Authors:** Leo Tomasevic, Hartwig Roman Siebner, Axel Thielscher, Fiore Manganelli, Giuseppe Pontillo, Raffaele Dubbioso

## Abstract

**Background:** The human primary sensory (S1) and primary motor (M1) hand areas feature high-frequency neuronal responses. Electrical nerve stimulation evokes high-frequency oscillations (HFO) at around 650 Hz in the contralateral S1. Likewise, transcranial magnetic stimulation (TMS) of M1 can evoke a series of descending volleys in the corticospinal pathway that can be detected non-invasively with a paired-pulse TMS protocol, called short interval intracortical facilitation (SICF). SICF features several peaks of facilitation of motor evoked potentials in contralateral hand muscles, which are separated by inter-peak intervals resembling HFO rhythmicity.

**Hypothesis:** In this study, we tested the hypothesis that the individual expressions of HFO and SICF are tightly related to each other and to the regional myelin content in the sensorimotor cortex.

**Methods:** In 24 healthy volunteers, we recorded HFO and SICF, and, in a subgroup of 20 participants, we mapped the cortical myelin content using the ratio between the T1- and T2-weighted MRI signal as read-out.

**Results:** The individual frequencies and magnitudes of HFO and SICF curves were tightly correlated: the intervals between the first and second peak of cortical HFO and SICF showed a positive linear relationship (r= 0.703, p< 0.001), while their amplitudes were inversely related (r= −0.613, p= 0.001). The rhythmicity, but not the magnitude of the high-frequency responses, was related to the cortical myelin content: the higher the cortical myelin content, the shorter the inter-peak intervals of HFO and SICF.

**Conclusion:** The results confirm a tight functional relationship between high-frequency responses in S1 (i.e., HFO) and M1 (i.e., as measured with SICF). They also establish a link between the degree of regional cortical myelination and the expression of high-frequency responses in the human sensorimotor cortex, giving further the opportunity to infer their generators.

## 1. Introduction

Peripheral sensory stimulation produces a complex set of regional responses in the contralateral somatosensory cortex, where afferent input is processed, and that can be detected by electroencephalography (EEG). For instance, electrical nerve stimulation at the wrist does not only evoke an early somatosensory evoked potential (SEP) at around 20 ms (N20) in the contralateral primary somatosensory area (S1) [1] but concurrently induces high-frequency oscillations (HFO) at about 650 Hz [1–5]. HFO are composed of two independent bursts, probably the first of subcortical (early-HFO) and the second of cortical origin (late-HFO), arising respectively before and after N20 [4,6]. Late-HFO are generated in S1, namely in Brodmann areas 3B (BA3B)[2] and 1 (BA1) [2], and are associated respectively to peaks from 1 to 3, and to peaks from 3 to 5 in the burst following N20 [7]. A study in monkeys employed concurrent extracellular single-unit recordings to show that the activity in BA3 is generated by bursts of 2 to 5 action potentials likely arising from pyramidal neurons, as suggested by the duration of the action potentials and their detectability with the EEG [8,9].

Phenomena that feature a rhythmic pattern at around 650 Hz can be elicited in the descending corticospinal tract with transcranial magnetic stimulation (TMS) of the human primary motor area (M1) [10–12]. This pattern of activity can be observed indirectly and non-invasively using a paired-pulse protocol. Indeed, if two TMS pulses are applied at interstimulus intervals ranging from 0.6 to 5 milliseconds and at an intensity around cortical motor threshold [13], or a subthreshold pulse after a suprathreshold one [14], it is possible to observe several peaks of facilitation of the motor evoked potentials (MEPs) at specific inter-stimulus intervals. The rhythmicity of these peaks of short interval intracortical facilitation (SICF) closely resembles the one of sensory evoked HFO in S1 or the rhythmicity of the descending I-waves evoked by TMS in the corticospinal tract [10,14–17]. While its exact neural origin is still debated, I-waves are thought to reflect the strength and temporal dynamics of cortical circuits that facilitate transsynaptic excitation of those pyramidal cells in M1 that send direct monosynaptic projections to the spinal motoneurons [16,18,19].

Given the cortical origin and shared rhythmicity of late HFO, on the one hand, and descending I-waves and the peaks of SICF, on the other hand, it has been hypothesized that these phenomena may be closely related [8,16]. Yet, studies that have directly examined whether and how their properties are related to each other, are lacking. Another question that remains to be addressed concerns the microstructural properties that determine the individual expression of HFO and I-waves in the sensorimotor cortices. One candidate is the degree of cortical myelination, because myelin determines crucial aspects of temporal processing in neuronal circuits, including conduction velocity, the synchronization of neuronal activity [20–22], and it is shaped by the firing rate of the action potentials [23,24]. Therefore, regional myelin content may sustain high-frequency phenomena and may set a limit on the maximal frequency that can be expressed in the cortical region.

In this study, we combined magnetic resonance imaging (MRI) of cortical myelination with recordings of HFO and SICF to test two hypotheses. We predicted that the individual expressions of high frequency activity in S1, as expressed by HFO, and high frequency activity in M1, as reflected by SICF, are tightly related in terms of rhythmicity and magnitude. We further hypothesized that the cortical myelin content in the sensorimotor cortex is related to the frequency or magnitude of the high frequency cortical activity, measured by HFO and SICF.

## 2. Methods

### 2.1 Subjects

Twenty-four subjects (14 female) aged 22–44 years (mean age ± SEM: 33.13 ± 2.64 years) were enrolled in the study after giving their written informed consent. The experiments conformed to the Declaration of Helsinki and were approved by the Ethics Committee of the University of Naples Federico II, Italy (N. 100/17). None of the subjects had a history of neurological disease or received drugs interacting with the central nervous system at the time of the experiment. All the subjects were right-handed according to the Edinburgh Handedness Inventory [25].

### 2.2 Experimental procedures

The experimental procedures are summarized in Fig.1. All participants underwent measurements of somatosensory HFO followed by paired-pulse TMS to obtain the individual SICF curve. Before starting with the TMS protocol, all subjects were screened for contraindications to TMS [26]. During the electrophysiology experiments, subjects were seated comfortably in a reclining chair with the right forearm placed in a prone position on the arm rest.

**Figure 1.**
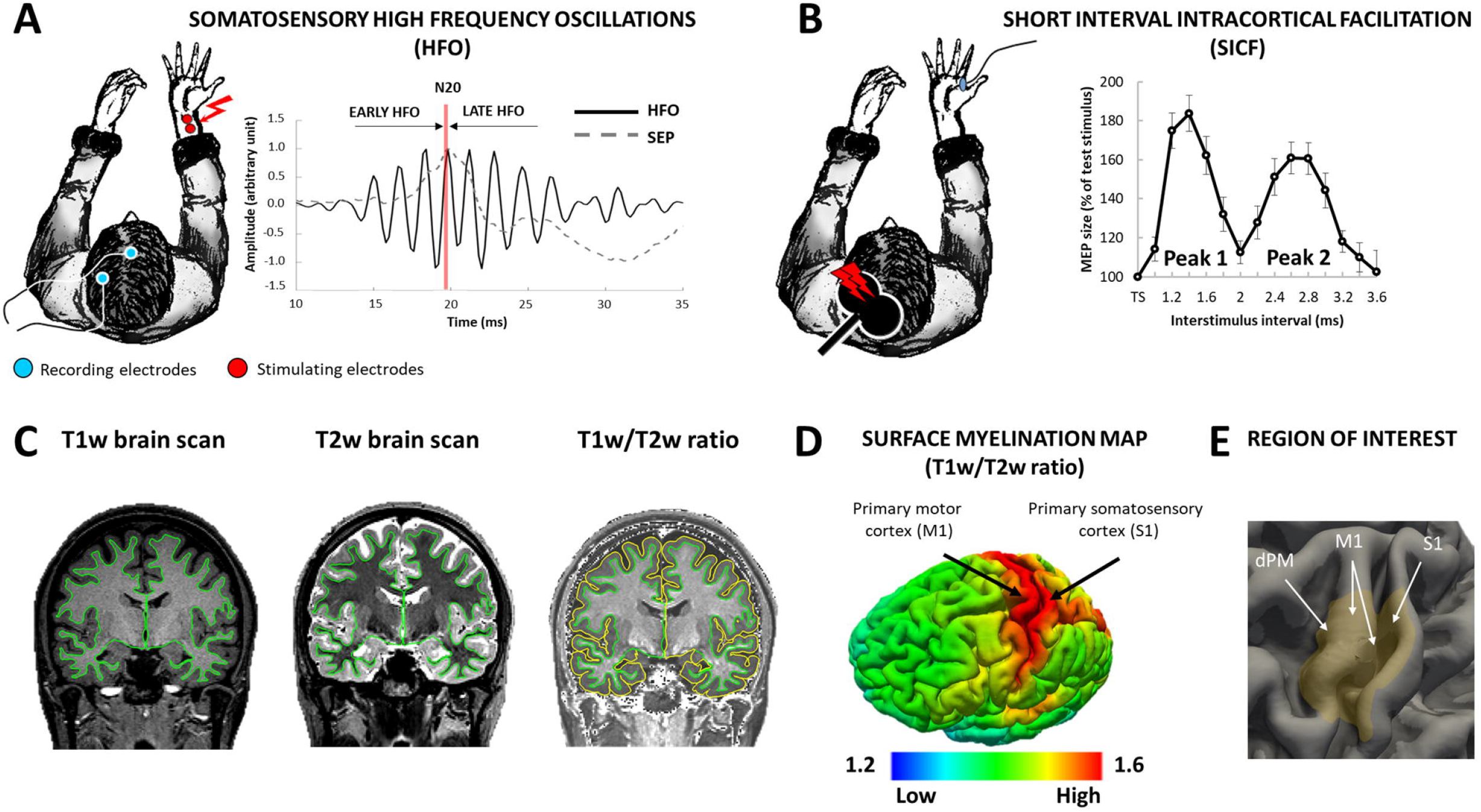
Experimental set-up. (A) Somatosensory evoked potential, SEP (dotted line) and high frequency oscillations, HFO (continuous line) after filtering and averaging EEG signal from CP3 active and Fz reference electrodes (light blue circles). HFO are divided in the early and the late components occurring before and after N20 peak respectively. (B) SICF curve over M1 at interstimulus intervals between 1.0 – 3.6 ms, covering the early facilitatory peaks (Peak 1 and Peak 2). (C) From left to right: T1w and T2w MRI scans with the white matter (green line) and cortical pial surfaces (yellow line); T1w/T2w ratios are estimated by dividing the T1w by the T2w image and the values are sampled on the mid-thickness surface, which is positioned halfway in-between the white matter surface and cortical pial surface. (D) Cortical myelination amount across different cortical regions. (E) Region of Interest used for surface analysis: dPM = dorsal Premotor cortex; M1 = primary motor cortex (crown and bank); S1 = primary somatosensory cortex.

### 2.3 Recordings of somatosensory cortical responses

Peripheral electrical stimulation of the right median nerve was performed at the right wrist placing the anode on the crease and the cathode 2 cm proximal. Three thousand monophasic square-wave pulses of 200 μs duration were delivered at a frequency of 5 Hz (Digitimer, Welwyn Garden City, UK). Stimulus intensity was adjusted to 120% of individual motor threshold, which is defined as the minimum stimulation intensity able to produce a small twitch of the abductor pollicis brevis muscle.

SEPs were recorded from scalp with Ag–AgCl cup electrodes placed at CP3 (active channel) and at Fz (reference), according to the international 10–20 EEG montage system [27]. We chose this montage because the bipolar parieto-frontal montage has been shown to be more sensitive to the BA3B component of HFO [28]. Cortical responses were recorded with a one-second-long trial at a sampling rate of 5 kHz using Signal software and CED 1401 hardware (Cambridge Electronic Design, Cambridge, UK). Pre-processing of the EEG data consisted in cubic-interpolation of the stimulus-related electric artefact from −0.2 to +6 ms [29] and exclusion of saturated trials. Subsequently, N20 peak latency and HFO were detected after band-passed filtering, 3-2000 Hz and 400-800 Hz respectively (zero-phase second order Butterworth filter) and averaging the trials (Fig. 1A). We used the N20 latency to distinguish the two, subcortical and cortical, bursts present in HFO.

Further HFO analysis was based on findings from the studies from Baker [8] and Telenczuk [9], which had shown that the two central and highest peaks represent cortical population spiking of BA3B, in line with Shimazu’s study [7]. Consequently, we selected, for the subcortical and cortical sources, the two highest peaks before and after N20 respectively, which happened to be the closest peaks to N20 and to be always positive in our montage. We did not consider the negative peaks and we put them to zero. This is for action potentials have a dominant polarity, and filtering of only positive or only negative peaks induces ringing effects. Our procedure yielded two HFO datasets that reflected the subcortical and cortical components and were comparable in shape to the SICF curve.

### 2.4 Assessment of short interval intracortical facilitation (SICF)

A biphasic TMS pulse was applied using a standard figure-of-eight coil (MC-B70 with outer diameter of each wing 97 mm) connected to a high-power magnetic stimulator (MagPro X100, Medtronic, Denmark). During the experiment the coil was held tangentially to the scalp with the handle pointing backwards and rotated away from the midline at 45 degrees. This way, the current induced in the brain was directed from posterior-lateral to anterior-medial, which is optimal for activating pyramidal neurons trans-synaptically via horizontal corticocortical connections [30]. We used biphasic stimulation since recent studies have consistently shown that SICF can be induced by a biphasic TMS pulse, while offering higher energy-efficiency than a monophasic TMS pulse and thus, enabling efficient stimulation at lower stimulus intensities [31–33].

The “hot spot” was defined as the scalp position over the left M1 where maximal motor evoked potentials (MEPs) was elicited in the contralateral first dorsal interosseous (FDI) muscle [34]. The signal was acquired via Ag–AgCl surface electrodes in a belly tendon montage, amplified, bandpass filtered (20 Hz-3 kHz) and digitized at a frequency of 5 kHz (Signal software and CED 1401 hardware, Cambridge Electronic Design, Cambridge, UK). We first determined the resting motor threshold (RMT), given in percentage of maximum stimulator output, as the minimum stimulus intensity that produced a MEP of 50 μV in the relaxed right FDI muscle in at least 5 of 10 trials [35]. Finally, MEP1mV was determined as the stimulus intensity, which elicited in the resting FDI a MEP of 1 mV on average in five consecutive trials.

The SICF curve was evaluated by applying paired-pulse TMS at fourteen inter-stimulus intervals (ISIs, Fig 1B), ranging from 1.0 to 3.6 ms and separated by steps of 0.2 ms. The intensity of the first stimulus was set to MEP1mV and the second stimulus at 90% RMT [14,36]. Fifteen stimuli for each ISI were tested in a randomized order. Mean MEP amplitude was calculated for each ISI and normalized to the MEP size elicited by the test stimulus alone. This yielded a SICF curve which covered the first and the second facilitatory peaks (Fig.1B).

### 2.5 Structural MRI

Participants underwent structural MRI of the brain immediately after the electrophysiological measurements had been completed. Three-dimensional MRI scans were acquired with a 3 T Trio Scanner equipped with an 8-channel head coil (Siemens Healthineers, Erlangen, Germany). The acquisition protocol included a T1-weighted volume acquired using a 3D Magnetization Prepared RApid Acquisition Gradient Echo (MPRAGE) sequence consisting of 192 sagittal slices with 1mm isotropic voxel resolution (TR/TE = 2300/2.96 ms; TI = 1100 ms; flip angle = 9°; matrix size = 256 x 240 x 192) and a T2-weighted volume using a 3D turbo spin echo sequence with 176 sagittal slices and 1mm3 voxel resolution (TR/TE = 3200/408 ms; flip angle = 120°; matrix size = 256 x 258 x176). MRI data of four subjects had to be excluded because of head movement related artefacts.

### 2.6 Surface-based analysis of cortical morphology: thickness, curvature, and myelination

Cortical reconstruction was performed with the FreeSurfer image analysis suite ver. 6.0.0 (http://surfer.nmr.mgh.harvard.edu/) [37]. The grey and white matter surfaces were defined by an automated brain segmentation process. If required, an experienced investigator (R.D.) manually corrected the automated segmentation, following the procedures outlined at https://surfer.nmr.mgh.harvard.edu/fswiki/Edits. The processes of surface extraction and inflation generated surface curvature, estimated from the mean curvature (or average of the principal curvatures) of the white matter surface [38] and cortical thickness, estimated at each point across the cortex by calculating the distance between the grey/white matter boundary and the cortical surface. Individual whole brain surface maps were smoothed with a 5 mm 2D Gaussian smoothing kernel [37] and the effect of surface curvature on cortical thickness was regressed out [39,40]. Individual curvature-corrected cortical thickness maps were registered to a common FreeSurfer template surface (fsaverage) for visualization and group analysis [41].

Procedures for mapping the myelin content of the cortical surface are described in detail by Glasser and Van Essen [42]. Briefly, a T1-weighted (T1w) and T2-weighted (T2w) image were registered to each other, and a ratio between the two images was calculated in each voxel of the brain, as a relative marker of myelin content (Fig. 1C). Specifically, the T1w image was brain-extracted and segmented into white matter and cerebro-spinal fluid using the FAST procedure [43] implemented in FSL [44]. The T2w image was brought into register with the T1w image using FSL flirt, employing a rigid registration with mutual information as cost function. At each voxel, the ratio between the normalized T1w image and the aligned normalized T2w image was calculated, which is believed to be contrast-sensitive to myelin [42]. To specifically examine only cortical voxels, the cortical ribbon was defined by voxels between white and pial surfaces, and the T1w/T2w ratios were sampled to each subject’s individual ‘native’ central cortical surface by using cubic interpolation in MATLAB. We regressed out curvature since curvature-associated modulations can obscure or distort myelination changes due to other variations in cytoarchitectonic [45]. We finally smoothed the data applying a 5 mm 2D Gaussian smoothing kernel [37]. The individual maps were registered onto the FreeSurfer group template for visualization and group analysis (Fig 1D).

## 3. Statistical analysis

For both SICF and HFO, we extracted values of frequency and amplitude that were then used for statistical analysis. Temporal centre of gravity (CoG) of each peak was calculated for the peak latency and the distance of the two latencies was used as frequency measure. Regarding the amplitude measure, we computed the sum of the area under the curve (AUC) [46] obtained from the first and second peak (not normalized data). In the case of HFO, we considered the two highest peaks before and after N20 for early- and late-HFO respectively.

Before applying parametric statistical tests, the normal distribution of all variables was verified by means of a Kolmogorov and Smirnov test. Then, as a preliminary step, we evaluated the effect of ISI on SICF considering raw and not normalized MEPs values. In detail, one-way repeated measure analysis of variance (ANOVA-rm) was conducted with the within subject factor ISI (14 levels for SICF: 1.0 - 3.6 ms in 0.2 ms steps). Post hoc twotailed one-sample t-tests were conducted to identify ISI with significant MEP change with respect to Test stimulus. In addition, paired t-tests were used to compare AUC of the first (AUCpeak1) and the second SICF peak (AUCpeak2) and regarding HFO, to compare the frequency and AUC of the early vs late component. The second part of the analysis consisted of:

i. Correlation of the temporal (i.e., frequency) and magnitude (i.e., AUC) properties of the early and late components of HFO with frequency and area of SICF peaks. Specifically, for the frequency analysis, we compared the distance between the CoGs of the HFO peaks with that obtained from SICF peaks. For spatial analysis, we correlated the amplitudes as the sum of areas that subtended the peaks’ curves for HFO and SICF.
ii. Correlation of the temporal (frequency) and magnitude (area) parameters of the early and late components of HFO, and of SICF with the individual estimates of cortical thickness (derived from T1-weighted MRI scans) and myelination (derived from T1w/T2w ratio mapping).

To visualize the spatial distribution of significant effects we computed surface-based analyses within the left sensory-motor cortex forming the hand knob. We defined our region-of-interest (RoI) as the cortical area covering the crown and the bank of M1 (BA4a and BA4p), the adjacent dorsal Premotor cortex (dPM, BA6) till the bottom of the precentral sulcus [40] and S1 (BA3 and part of BA1) till the top of the postcentral gyrus crown (Fig 1E). The multimodal parcellation atlas derived from the Human Connectome Project [39] was used to visualize the border between premotor, motor, and somatosensory areas. In addition, we identified the left primary visual cortex (V1, BA17) and the left secondary visual cortex (V2, BA18) as “control” RoI to demonstrate the spatial specificity of our findings.

To visualize the spatial distribution of significant effects we computed surface-based analyses within the RoI on the FsAverage template by using Freesurfer software [41]. These analyses were performed vertex-wise, followed by cluster-wise corrections for multiple comparisons based on permutation method [47](cluster-determining threshold: p<0.01, cluster-wise p<0.05) [41]. Age at the time of MRI and sex were included in the model as nuisance variables.

The alpha inflation due to multiple comparisons was faced according to the Bonferroni procedure.

Descriptive statistics are reported as mean ± standard error of the mean (SEM). All statistical analyses used IBM SPSS Statistics software (Version 22 for Windows, New York City, USA).

## 4. Results

### 4.1 Short-latency intracortical facilitation (SICF)

The paired-pulse measurements showed consistent SICF in all individuals. SICF measurements yielded a main effect of ISI on mean MEP amplitude (F_(4.419, 101.644)_= 8.393, p < 0.001). Mean MEP amplitude elicited by paired-pulse stimulation were larger at ISIs of 1.2-1.8 (peak1) and 2.2–3.0 ms (peak2) relative to single-pulse TMS (all paired t-tests: p<0.033). These peaks were separated by a trough at ISI of 2 ms, where paired stimulation had no significant effect on MEP size (paired t-test: p= 0.724). The magnitude of these two peaks was expressed as AUC, the AUC of the first peak was significantly higher with respect to the second one (AUCpeak1= 2.14 ± 0.15 vs AUCpeak2= 1.88 ± 0.14, p= 0.011). The mean ± SD CoG of peak1 and peak2 were 1.48 ± 0.08 ms and 2.63 ± 0.16 ms respectively, with an interpeak interval of 1.16 ± 0.14 ms.

### 4.2 High-frequency cortical oscillations (HFO)

Subcortical and cortical HFO did not differ regarding peak-to-peak distance (p= 0.873): 1.54 ± 0.02 ms (early-HFO) vs 1.55 ± 0.06 ms (late-HFO), nor regarding the amplitude (p= 0.511): 0.35 ± 0.04 μV^2^ x 10^-3^ (early-HFO) vs 0.33 ± 0.04 μV^2^ x 10^-3^ (late-HFO).

### 4.3 Relationship between SICF and HFO

The individual expressions of late-HFO and SICF were tightly correlated (Fig. 2). The individual frequencies of HFO and SICF, i.e., the interval between the first and second peak of cortical late-HFO and SICF, showed a positive linear relationship (r= 0.703, p< 0.001, Fig. 2A). Conversely, the response amplitudes of cortical late-HFOs and SICF, indexed by the area under the curve, showed a negative linear relation (r= −0.613, p= 0.001, Fig. 2B). No significant correlation was observed between the frequencies (r= 0.016, p= 0.941) and amplitudes (r= - 0.394, p= 0.057) of sub-cortical early-HFOs and SICF. Significance level was set at p< 0.006 after correction for multiple comparisons.

**Figure 2.**
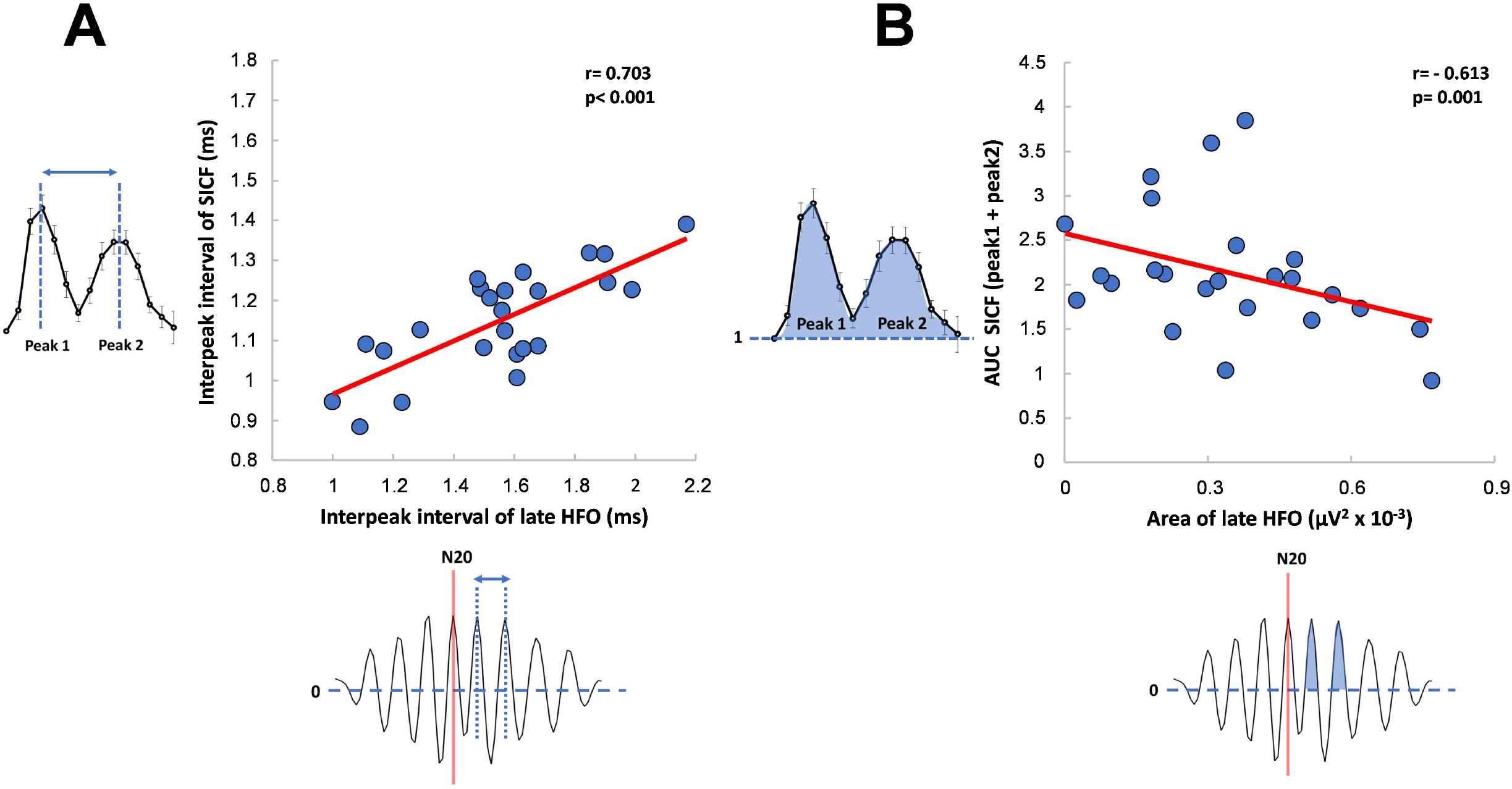
Relationship between somatosensory cortical high frequency oscillation (late HFO) and SICF. (A) Significant positive correlation between the interpeak interval of late-HFO and that elicited by SICF. (B) Significant negative correlation between the Area under the curve (AUC) of the first and second SICF peak and the AUC of the late component of HFO. Significant correlations are indicated in bold and by continuous lines. Significant p value <0.006.

### 4.4 Relationship between cortical myelination and cortical high-frequency responses patterns

The individual peak-to-peak intervals of both SICF and cortical late-HFOs showed voxel-based correlation with the myelin content in the S1 and with the crown of precentral gyrus (Fig. 3A and 3C). After cluster-based multiple comparison correction, surface-based correlation analyses revealed that the T1w/T2w ratio correlated with the individual frequency of cortical HFO in a cluster located posterior to the central sulcus, in the S1 (Fig. 3A; peak correlation at x, y, z = −35.3, −30, 52.8). For SICF, the correlation between the peak-to-peak interval of SICF and cortical myelin content was significant in a cluster covering mainly S1 along the whole hand knob (Fig. 3C; peak correlation at x, y, z = −50.3, −17.8, 51.2). Correlation analysis did not disclose any significant cluster in S1 or M1 that displayed a relation between the degree of cortical myelination and the amplitude of late-HFO or SICF. As for other cases, the subcortical early-HFO did not show significant correlation with myelin content. We also found no significant cluster when testing for a linear relationship between cortical thickness of the left S1 or M1 and the temporal and magnitude properties of early-HFO, late-HFO and SICF. Importantly, in the control RoIs, V1 and V2, we did not find any significant cluster when correlating cortical myelination and late-HFO or SICF.

**Figure 3.**
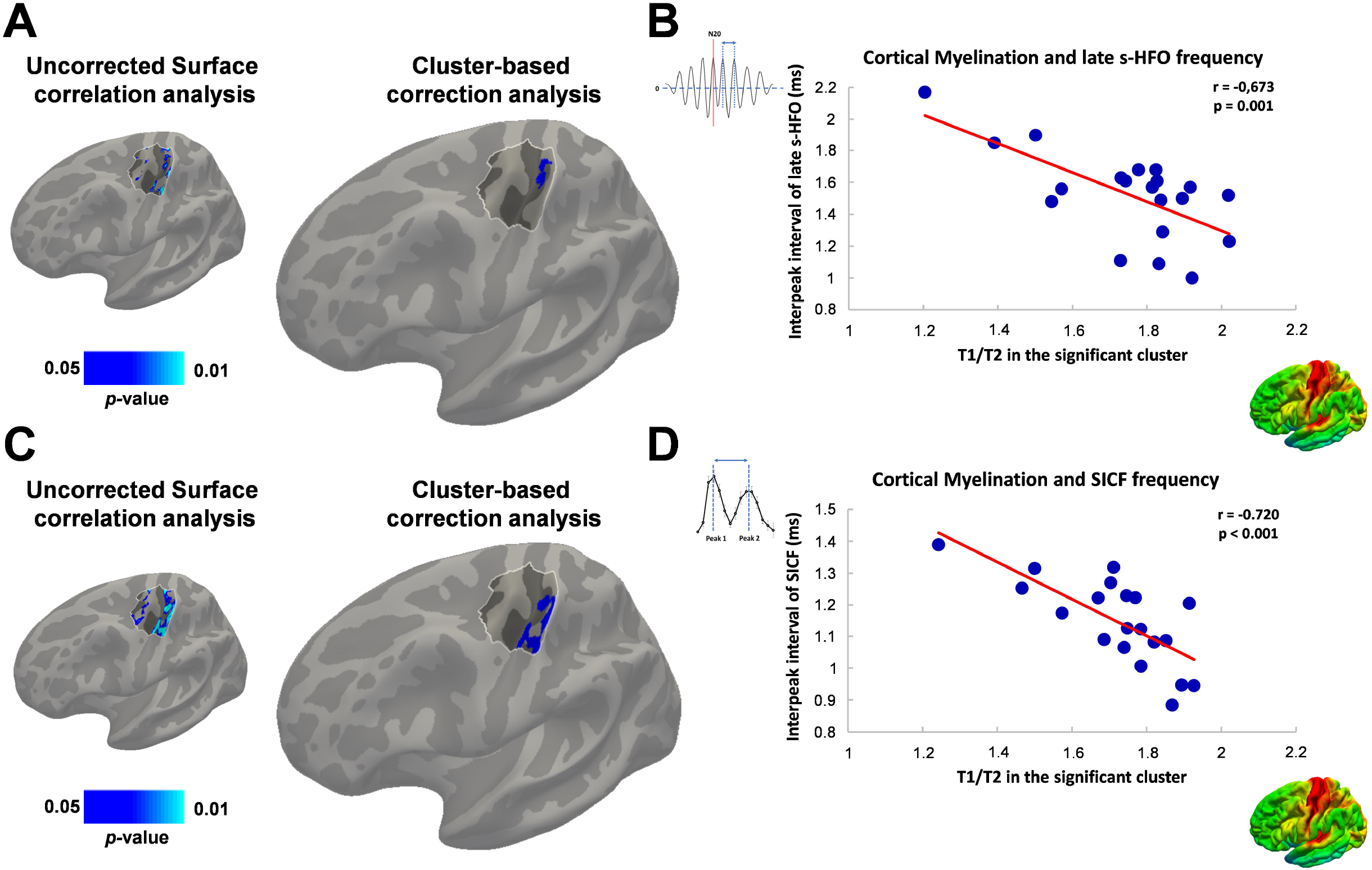
Relationship between cortical myelination and somatosensory high frequency oscillation (late-HFO) and SICF. (A) Surface-rendered statistical parametric map shows voxels with a negative linear relationship between cortical myelination and the inter-peak interval of late-HFO (uncorrected p-value < 0.05, left panel). The significant cluster is located in the primary somatosensory cortex (BA3) of the hand knob region peaking at x-,y-,z-coordinates −35, −30, 53 (cluster-wise corrected, p= 0.039, right panel). (B) Linear correlation between the T1w/T2w estimated myelination in the significant cluster and the interpeak interval of the late-HFO. (C) Surface-rendered statistical parametric maps show voxels with a negative relationship between cortical myelination and the inter-peak interval of SICF mainly in the primary somatosensory cortex and the crown of precentral gyrus (uncorrected p-value < 0.05, left panel). The significant cluster is located at the hand knob region in the primary somatosensory cortex region peaking at x-,y-,z-coordinates −50, −18, 51 (cluster-wise corrected, p= 0.006). (D) Linear correlation between the T1w/T2w estimated myelination in the significant cluster and the interpeak interval of the SICF. The region of interest (ROI) is highlighted in all brain maps.

## 5. Discussion

We found that the individual frequencies and magnitudes of late-HFO and SICF curves were tightly correlated. The intervals between the first and second peak of cortical HFO and SICF showed a positive linear relationship, while their magnitudes had a negative relation. The higher the cortical myelin content was, the shorter the interpeak intervals were of late-HFO and SICF, but there was no relation between amplitudes and myelin. Early-HFO, that are of subcortical origin and used in our study as a control condition, were not correlated either with SICF or with myelin content in the cortex.

### 5.1 High-frequency responses in the S1 and M1 are mutually related

The high-frequency response in the S1 evoked by peripheral electrical stimulation (i.e., late-HFO) was positively related to the rhythmicity evoked in M1 with TMS (Fig 2A). The generators of late-HFO activity are bursting pyramidal neurons in BA3B [8,9], most likely the thick tufted pyramidal neurons from layer 5b that are known to be activated by passive touch [48,49]. In fact, not only they are intrinsic bursting neurons that exhibit a bursting frequency compatible with HFO, but they also have adequate structure and geometry to be detected by EEG: they are well aligned in a parallel fashion with dendrites spreading from the lower part of layer 5 up to layer 1 on the cortical surface [50–52]. Thick tufted neurons are pyramidal-tract neurons which have cortico-subcortical, mainly cortico-spinal axonal projections [53,54]. Interestingly, pyramidal-tract neurons display high-frequency descending axonal activity in response to cortical stimulation, defined as indirect waves (I-waves), as observed after local cortical stimulation of the sensory and the motor primary areas [15].

Under the premise that the SICF curve is a non-invasive reflection of I-waves [13,14], our findings suggest a tight relationship between the burst generation of thick-tufted pyramidal-tract neurons in the sensory and those in the motor area.

Lastly, invasive epidural recordings in humans have shown that biphasic transcranial stimulation at low stimulation intensity can also evoke responses at around 330 Hz, which are overridden by I-waves at higher intensities (Fig 2, [12]). Responses at 330 Hz were also observed in animals [55], but again with invasive recordings. Although, our data are not suited to clarify how 330 Hz responses impact on SICF measures, and we cannot completely exclude their influence on the estimation of SICF magnitude or peak latencies, we do not think they had a major impact, apart from adding noise to our measurements.

### 5.2 Inverse relation of the magnitude of sensory and motor high-frequency responses

The individual magnitudes of late-HFO and SICF curves showed an inverse relationship of areas underlying the peaks of the responses (Fig 2B). Pharmacological and TMS studies showed that both protocols are dependent on GABA-ergic inhibitory circuits. Specifically, it has been suggested that late-HFO reflect the activity of inhibitory interneurons that produce feedforward inhibition of pyramidal neurons [4], and analogously the high frequency repetitive discharge of the corticospinal axons in I-waves is the result of recruitment of GABA(A)-ergic interneurons together with highly synchronized excitatory neurons [16]. Pharmacological studies showed increased SICF when a higher presynaptic release of glutamate was induced [56] and the facilitation was inhibited by positive allosteric modulators of GABA(A) receptor [57,58], but not GABA(B) agonist [57]. Importantly, these modulations are related only to the later I-wave peaks. Similarly, GABA(A)-receptor antagonist, bicuculline methiodide, increased the number of peaks of late-HFO [59]. This fits again with the characteristics of intrinsic bursting thick tufted pyramidal-tract neurons. In fact, GABA(A) inhibitory interneurons [60] can promote pyramidal neurons from single spike to bursting neurons [61,62], keeping the first peak stable and influencing only later components of the burst.

Since our study was not designed to clarify the mechanism behind the inverse relationship between the late HFO response generated in S1 and the amplitude of SICF peaks in M1, we can only speculate about the neural underpinnings. Based on the existing literature, we would like to put forward three hypotheses which are not mutually exclusive.

i. Firstly, HFO and SICF amplitudes may evidence reciprocal interaction between S1 and M1, resulting in a stronger inhibition of cortical neurons producing late HFO in the presence of stronger activity of cortical neurons responsible for facilitation peaks in SICF and vice versa. Accordingly, mean MEP amplitudes at rest have been reported to be higher when oscillatory activity in the hand knob in the alpha band (mu-activity) is increased [63–65], the latter representing a state of reduced neural excitability and activity of the somatosensory cortex [66].
ii. Secondly, the two different types of stimulation, peripheral nerve stimulation versus transcranial motor cortex stimulation, may have opposite effects on intracortical GABA-ergic circuits impinging on pyramidal tract neurons. While peripheral afferent input reaches the pyramidal neurons by thalamic projections to layer 2/3, 4 and 5 [67], TMS targeting the motor hand representation may primarily excite dorsal premotor sites in the crown-lip region of the precentral gyrus, which then excite the M1 via short-range cortico-cortical connections [40,68,69]. Moreover, simulations showed that TMS activates not only pyramidal neurons but also basket cells [69].
iii. Thirdly, the origin of neuronal activity probed with the two techniques differs. While SICF is the result of propagation of action potentials at the cortical and corticospinal level, EEG records neuronal activity expressed by polarization of the dendritic tree of pyramidal cells after action potential backpropagation. Here it is important to point out that larger action potentials at the soma are associated with a higher attenuation of the signal backpropagating in the dendrites [70].

In any case, this finding warrants further study, as this would advance our understanding of sensorimotor interaction in the human cortex.

### 5.3 The frequency of high-frequency cortical responses is related to cortical myelin content

MRI-based cortical myelin mapping identified cortical clusters in S1 where regional myelin content correlated positively with the individual frequency of late-HFO and SICF peaks (Fig 3). The higher the frequencies, the stronger was regional axonal myelination. This finding is compatible with the notion that the ability to generate faster rhythmic activity requires a higher myelin content because less myelinated axons show more jittered transmission of action potentials [22]. A high degree of axonal myelination secures a high level of synchrony that is necessary to allow precise timing of action potentials’ encounter at the postsynaptic neuron, to activate postsynaptic signal propagation [21]. This applies to bursting activity as well. High level of synchrony is needed to preserve the temporally precise forward transmission in down-stream neurons of all the spikes present in a burst, as is the case for I-waves. In fact, two consecutive spikes delivered by a multitude of neurons would merge at the postsynaptic level, if both their temporal distance and spiking synchrony were low (Fig. 4). Moreover, large diameter and myelinated axons are not only more suitable for temporally precise transmission, but they are also more resilient to high frequency spiking [71]. Finally, our data suggest that myelin content might be responsible for the frequency correlation between HFO and I-waves, but it also implies that myelin content of pyramidal tract neurons in S1 and M1 are mutually related.

**Figure 4.**
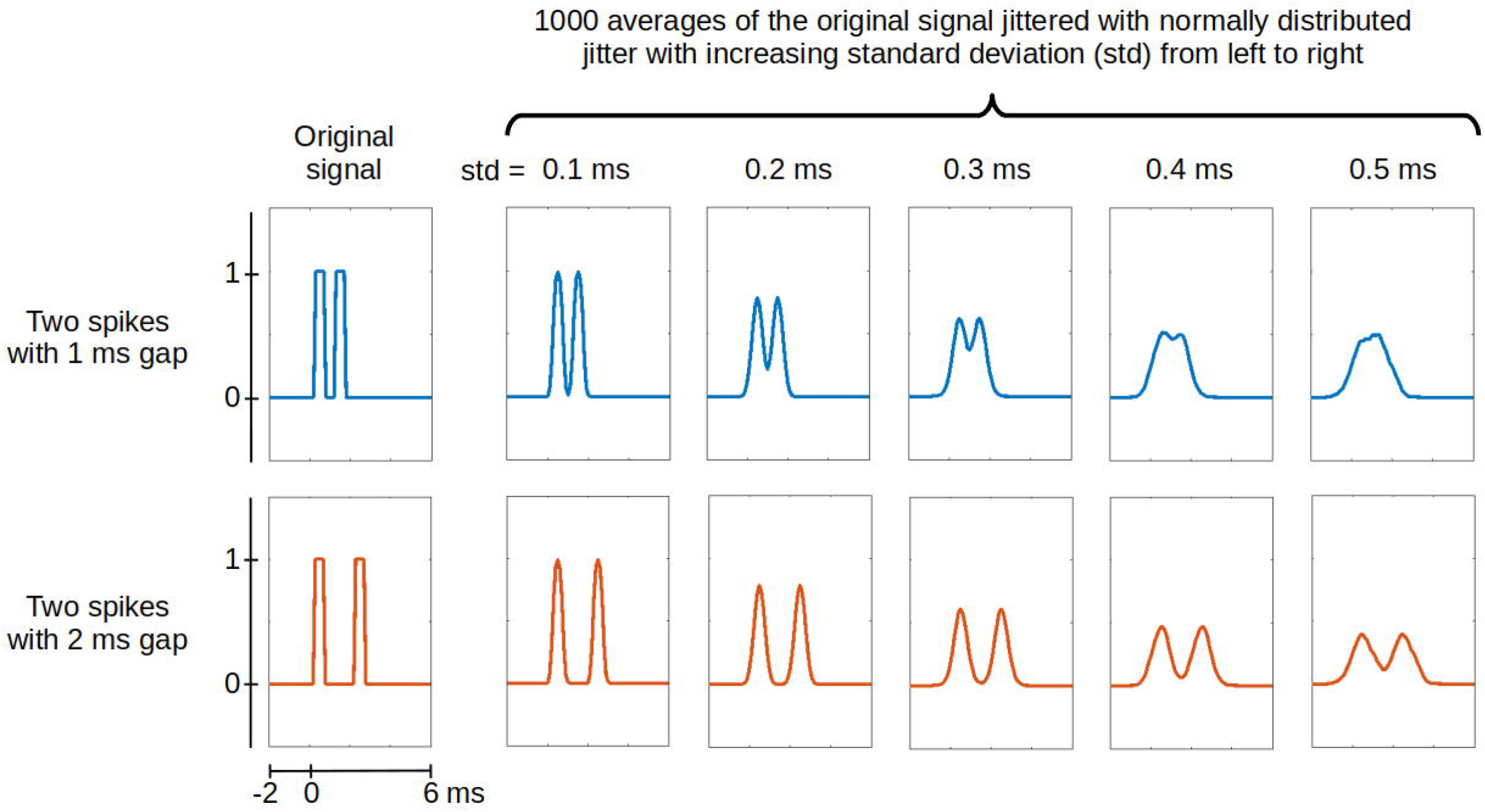
Simulation of the arrival of bursts at the postsynaptic neuron. We used a combination of two rectangular functions to mimic a burst of two action potentials. In the first row, there are two spikes with an inter-spike gap of 1 ms, while the gap is 2 ms in the second row. The first column shows the original signals, that are averaged 1000 times in the following columns after being jittered with a normally distributed jitter. In each column the jitter was increased by increasing the standard deviation of its distribution. The simulation shows that a slower burst is more resistant to lack of synchrony when encountering other bursts at the postsynaptic neuron.

We did not find a correlation between the myelin content and magnitude of high-frequency activity, neither in M1 nor in S1. We hypothesized that a higher synchrony would involve a higher level of overlap of action potentials at postsynaptic neurons providing higher peaks in both HFO and SICF. But, as discussed in the previous paragraph (5.2), the amplitudes are sensitive to GABA-ergic modulation, suggesting that the overall number of activated neurons has a higher impact on the magnitude than the synchrony among them [57,58]. It may even be possible that there is a negative relationship between magnitude and frequency, since a loss of myelin may increase neuronal excitability [72]. This consideration may be relevant in pathological states associated with a marked loss of cortical myelin, and this possibility is worth investigating in future studies on patients with cortical dysfunction and demyelination.

We also expected to find that clusters in M1 would show a positive relationship between regional myelination and I-wave rhythmicity, since I-waves are generated in the motor cortex [15], but significant clusters were confined to the sensory cortex in the postcentral gyrus. The voxel-based correlation map (Fig. 3) also showed spots on the crown of the precentral gyrus, where TMS produces the strongest electrical field [40,73], but the correlation didn’t survive after cluster-based multiple comparisons. This might be due to the structural inhomogeneity of the motor cortex, where, contrary to other brain regions, layer 5b also contains large Betz cells [74], which likely introduce inter-individual variability in the estimation of regional myelin related to the high frequency activity. In fact, Betz cells display a different shape, dendrites, and size compared to the surrounding pyramidal neurons [75]. In addition, they are not distributed homogeneously in the precentral gyrus, as they show a mediolateral gradient with a cluster in the precentral hand knob, the region of our interest [76]. Therefore, a larger sample size may be needed to reveal a significant correlation between regional myelin content in precentral M1 and the individual frequency of late-HFO and SICF peaks. This issue remains to be addressed in more detail in future studies.

## 6. Conclusions

Our study sheds new light on high-frequency responses in the primary sensory and motor cortices in humans. We showed a strong link between late-HFO and I-waves represented by SICF. Both cortical phenomena may reflect the activity of intrinsic bursts of thick tufted pyramidal-tract neurons that are generated in strong functional interaction with GABA-ergic inhibitory interneurons. Our results also suggest a strong link between the rhythmicity of the cortical high-frequency responses and regional myelin content, underscoring the importance of regional myelination for the precise timing of neuronal activity in the human cortex. The results of this study also have important implications for the clinical neurophysiological assessment of cortical dysfunction. For instance, abnormally enlarged high-frequency somatosensory oscillations were shown in patients with movement disorders, such as Parkinson disease and myoclonus epilepsy [77,78], and slowing of high-frequency bursts was observed in myoclonus epilepsy [77] and multiple sclerosis [79], the latter being also affected by the reduction of cortical myelin content [80,81].

## CRediT authorship contribution statement

**Leo Tomasevic:** Conceptualization, Visualization, Software, Project administration, Writing-original draft, Writing-Review & editing.

**Hartwig Roman Siebner:** Visualization, Writing-original draft, Writing-Review & editing.

**Axel Thielscher:** Methodology, Software.

**Giuseppe Pontillo:** Data curation, Methodology, Writing-original draft.

**Fiore Manganelli:** Visualization, Writing-original draft.

**Raffaele Dubbioso:** Data curation, Methodology, Formal analysis, Investigation, Visualization, Writing-original draft, Writing-Review & editing.

## Abbreviations

AMT: Active Motor Threshold
M1: Primary Motor Cortex
HFO: High Frequency Oscillations
RMT: Resting Motor Threshold
S1: Primary Somatosensory Cortex
SICF: Short Interval Intracortical Facilitation
SEP: Somatosensory Evoked Potential
TMS: Transcranial Magnetic Stimulation

